# metaVaR: introducing metavariant species models for reference-free metagenomic-based population genomics

**DOI:** 10.1101/2020.01.30.924381

**Authors:** Romuald Laso-Jadart, Christophe Ambroise, Pierre Peterlongo, Mohammed-Amin Madoui

## Abstract

**Motivation:** The availability of large metagenomic data offers great opportunities for the population genomic analysis of uncultured organisms, especially for small eukaryotes that represent an important part of the unexplored biosphere while playing a key ecological role. However, the majority of these species lacks reference genome or transcriptome which constitutes a technical barrier for classical population genomic analyses.

**Results:** We introduce the metavariant species (MVS) model, a representation of the species only by intra-species nucleotide polymorphism. We designed a method combining reference-free variant calling, multiple density-based clustering and maximum weighted independent set algorithms to cluster intra-species variant into MVS directly from multisample metagenomic raw reads without reference genome or reads assembly. The frequencies of the MVS variants are then used to compute population genomic statistics such as *F*_*ST*_ in order to estimate genomic differentiation between populations and to identify loci under natural selection. The MVSs construction was tested on simulated and real metagenomic data. MVs showed the required quality for robust population genomics and allowed an accurate estimation of genomic differentiation (Δ*F_ST_ <* 0.0001 and < 0.03 on simulated and real data respectively). Loci predicted under natural selection on real data were all found by MVSs. MVSs represent a new paradigm that may simplify and enhance holistic approaches for population genomics and evolution of microorganisms.

**Availability:** The method was implemented in a R package, *metaVaR*. https://github.com/madoui/MetaVaR

## 1 Introduction

Thanks to progresses in sequencing technologies and metagenomics, microorganism genomic resources increased considerably the last two decades allowing a better understanding of microbial ecology through the analysis of community assemblies that contains a large number of uncultured species (Chariton, 2019). This is especially the case for marine, soil and gut microbiomes that were intensively investigated thanks to large sequencing consortia like *Tara* Oceans (Carradec *et al.*, 2018; Ibarbalz *et al.*, 2019), TerraGenome (Vogel *et al.*, 2009) or MetaHit (Ehrlich *et al.*, 2011).

Currently, to address questions relative to molecular evolution and population genomics of uncultured species, the efficient use of whole-genome metagenomic data requires reference genome or transcriptome sequences *a priori* selected for target species. Typically metagenomic reads are aligned on reference sequences for variant calling followed by the computation of population alleles or amino acids frequencies and derived population genomics metrics. This approach was applied notably in gut microbiome of vertebrates (Schloissnig *et al.*, 2013; Garud and Pollard, 2020) and invertebrates (Ellegaard and Engel, 2019), marine bacteria (Delmont *et al.*, 2019; Salazar *et al.*, 2019) or crustaceans (Madoui *et al.*, 2017).

In the context of metagenomics, the reads alignment filtering step is critical to avoid cross-mapping of reads from a given species to the reference genome of another species. The main filters are, first the selection of genomic regions with a depth of coverage within an expected range specific to the species abundance and second, the minimum identity percentage of a read aligned to the reference to be considered as belonging to the targeted species (Costea *et al.*, 2017; Madoui *et al.*, 2017).

To increase the number of reference sequences of organisms found in environmental samples, several methods were developed to produce metagenome assembled species (MAGs) from whole-genome metagenomic sequencing. These approaches were successfully applied on prokaryotes (Sedlar *et al.*, 2016; Parks *et al.*, 2017; Stewart *et al.*, 2019) but the main limit of the alignment-based approach still relies on the availability, completeness and quality of reference genomes that can be constructed from metagenomic data (Parks *et al.*, 2017). The approach took recently advantages of long-read sequencing (Huson *et al.*, 2018; Somerville *et al.*, 2019). However, due to the large genome size of eukaryotes and the difficulties to obtain high molecular weight DNA, yet, such approaches did not produce results on eukaryotes.

To identify nucleic variants from metagenomic data, an alternative to the alignment-based approach has been recently proposed through reference-free variant calling based on the *DiscoSNP++* tool (Uricaru *et al.*, 2015). This software detects the variants by identifying bubbles in a *de Bruijn* graph built directly from the raw metagenomic reads. Then, the variants can be relocated on genomes of interest if available. Comparing to alignment-based variant calling applied on metagenomic data, *DiscoSNP++* was presented as less sensitive but more specific in term of recall, and more accurate in term of allele frequency, especially in noncoding regions (Arif *et al.*, 2018). As population genomic analysis is very sensitive to the accuracy of the allele frequencies, we can consider that the use of *DiscoSNP++* could be preferable to alignment-based approach for population genomics based on metagenomic data.

In this study, we introduce the notion of metavariants, i.e variants detected directly from raw metagenomic reads without reference genome. We propose an effective method to cluster the metavariants by species and propose these clusters as a new species representation named metavariant species or MVS. We establish a formal definition of the metavariant and MVS. The relevance of the MVSs was tested on simulated and real metagenomic data. The clustering method we developed to build the MVSs was implemented in a *R* package called *metaVaR* and compared to state-of-the-art clustering algorithms.

## 2 Methods

### 2.1 From metavariants to metavariant species

#### 2.1.1 Variable loci and metavariants

We define a metavariant as a single nucleotide variant detected directly from metagenomic data without reference genome. We use metavariants produced by *DiscoSNP++* and consider only metavariants located on loci producing one metavariant. Due to the absence of reference genome, the reference (*a*) and alternative (*b*) nucleotides are chosen by *DiscoSNP++* based on the alphabetic order. In a single sample, *a* and *b*, are characterized by the count of reads supporting them and a locus *l* that harbors a metavariant, can be represented by its depth of coverage *c* as the sum of reads supporting *a* and *b*. Each locus *l* is described by the *m* sample supporting counts *l* = {*c*_1_, …, *c*_*m*_}. The *n* metavariant loci row vectors *l*_*i*_ generated from *m* samples metagenomic data are gathered in the *n* * *m* depth of coverage matrix, *L* = {*l*_*i*_} = (*c*_*ij*_) ∈ ℕ^*n*\**m*^, *i* ∈ {1, …, *n*}, *j* ∈ {1, …, *m*}.

#### 2.1.2 Clustering of metavariants into metavariant species

A metavariant species or MVS corresponds to a set of intra-species metavariants of the same species. We suppose that *L* contains both inter and intra-species metavariants, thus MVSs can be represented by pairwise disjoint subsets of *L* not covering *L*.

As for metagenomic contigs binning (Sangwan *et al.*, 2016), we consider that the depth of coverage of the variable loci of the same species covariates across samples and this constitutes a species signature. Thus, MVSs can be found by clustering *L* based on its values. However, the complexity of metagenomic data rises several issues. First, the number of species and corresponding MVS is unknown. Second, the initial set of metavariants contains an admixture of inter and intra-species metavariants coming from entire genomes including repeated regions. Only intra-species metavariants from single copy loci are informative for population genomics. Third, the genome size and the polymorphism rate highly vary between species. This impacts the loci depth of coverage and the number of variants by species.

#### 2.1.3 Metavariant species construction steps and algorithms

To create MVSs, we propose the following approach, described in details in the following sections:

1. Reference-free metavariant calling with *DiscoSNP++* from raw metagenomic data (figure 1.B).
2. Metavariants filtering and construction of the loci depth of coverage and metavariant frequencies matrices (figure 1.B).
3. Multiple density-based clustering (*mDBSCAN*) of the metavariants. Each clustering generates a set of disjoint metavariant clusters (*mvc*) (figure 1.C).
4. Each *mvc* is scored according to its size and expected depth of coverage distribution of its loci (figure 1.D).
5. A *maximum weighted independent set* (WMIN) algorithm is applied on all *mvc* to select a subset of *mvc* as potential MVSs (figure 1.E).
6. Selection of the metavariant loci based on coverage expectation for robust population genomic analysis (figure 1.F).
7. Computation of population genomic metrics for each MVS (figure1.G).

**Fig. 1.**
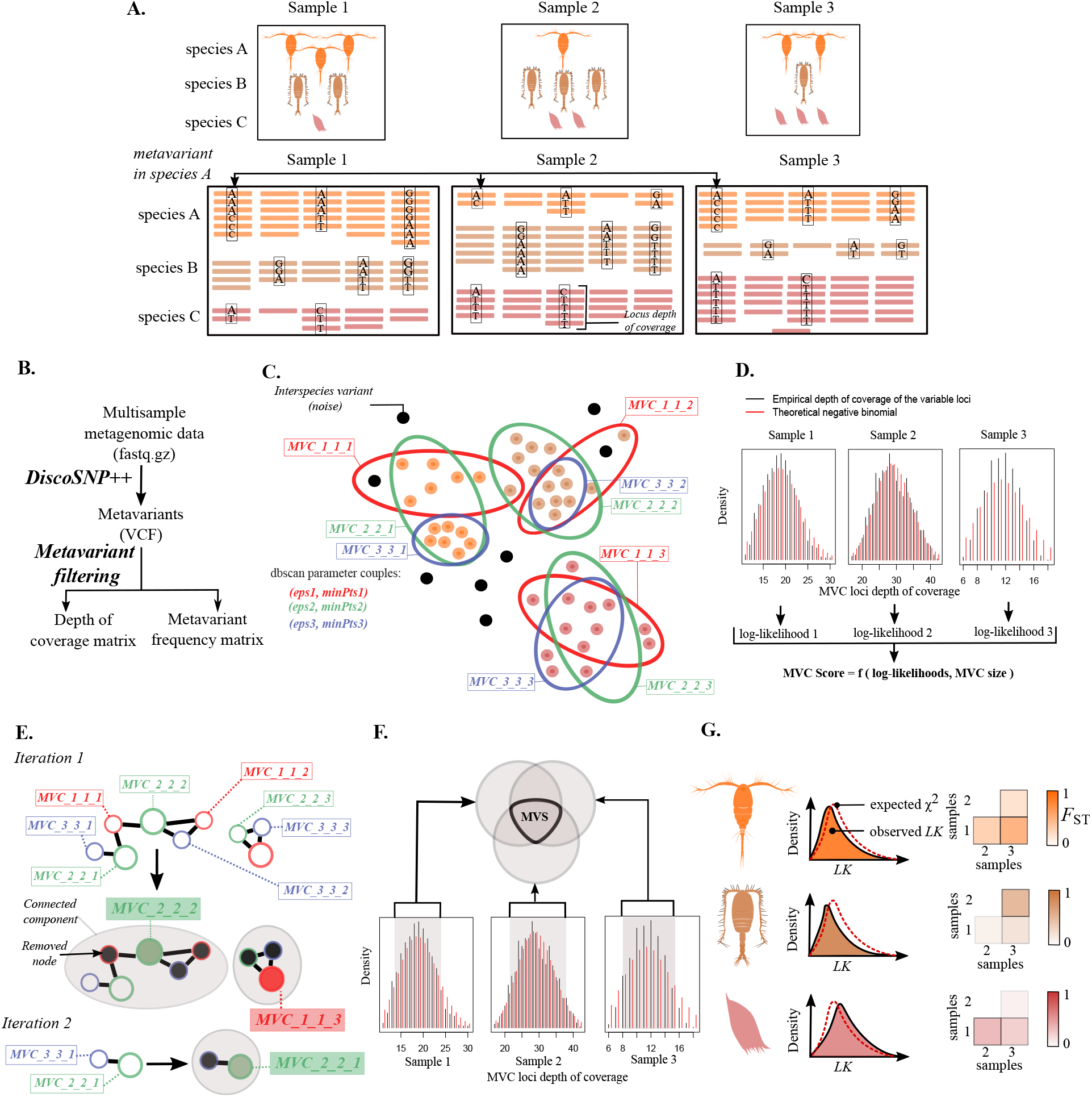
Metavariants species construction from metagenomic data with metaVaR. A. Environmental samples and metagenomic high-throughput sequencing. The example contains three different species. Species A generates orange reads, species B brown reads and species C red reads. B. Variant calling from raw metagenomic data. C. Multiple density-based clustering of metavariants mDBSCAN-WMIN. Black points represent inter-species variant while other colored points refer to intra-species variants. Circle colors represent the dbscan parameters. D. Metavariant cluster scoring. E. Maximum Weighted Independent Set algorithm. Each node is a cluster of metavariants, their circle color represent the dbscan parameters used to built the cluster. Grey zones represent the connected components. Colored nodes are MWIS and black nodes are MWIS neighbors.F. Metavariant filtering for MVS construction. Grey zones correspond to the single-copy loci. G. Population genomics of MVS

#### 2.1.4 Multiple density-based clustering of metavariants

To cluster *L*, we used density-based clustering (*dbscan*) (Campello *et al.*, 2013), a clustering algorithm that requires two input parameters: epsilon (*ϵ*) and minimum points (*p*) corresponding respectively to the minimum distance between two points to be considered as members of the same cluster and the minimum number of points to extend a cluster. Given *ϵ* and *p*, *dbscan* generates a disjoint set of metavariant clusters {*mvc*_*ϵ,p*_ ⊂ *L*}. Intuitively, for *L* generated from real data, there might be no optimal (*ϵ, p*) enabling the best reconstruction of the clusters according some criteron. Instead of choosing arbitrary one (*ϵ, p*) couple, we run *dbscan* using a grid of (*ϵ, p*) values. This multiple density-based clustering (mDBSCAN) produces a set of possibly overlapping *mvc*. We called this set *MVC* which is restricted to *mvc* containing more than 1,000 metavariants by default.

#### 2.1.5 Metavariant clusters scoring

Several metagenomic binning approaches use Gaussian or Poisson distributions to model genome sequencing coverage (Sedlar *et al.*, 2016). However, due to its overdispersion, the sequencing depth of coverage can be better approximated using a negative binomial (NB) distribution (Robinson and Smyth, 2007). Let *mvc*_*ϵ,p,k*_ ∈ *MVC* denotes the *k*^th^ *mvc* computed with parameters (*ϵ, p*). For each *mvc*_*ϵ,p,k*_, in each sample, we compare the observed and expected NB coverage distribution of the loci using *fitdistrplus* (L. and Dutang, 2015). For all possible *mvc*_*ϵ,p,k*_, we compute *d*_*ϵ,p,k*_, the mean across *m* samples of the log-likelihood of the fitting with *θ*_*j*_, the negative binomial distribution parameters in sample *j*.

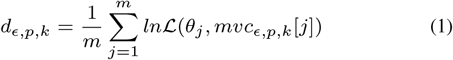

with *mvc*_*ϵ,p,k*_ [*j*] being the depth of coverage of the metavariant in the *j*^th^ sample in *mvc*_*ϵ,p,k*_. We also compute *d̅*_*ϵ,p,k*_ ∈ [0, 1], the corrected mean log-likelihood of each cluster,

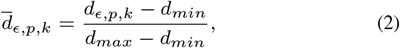

with *d*_*min*_ and *d*_*max*_ being respectively the smallest and the highest mean log-likelihood observed over all *mvc* ∈ *MVC*. We also correct the size *s*_*ϵ,p,k*_ of each cluster such as *s̅*_*ϵ,p,k*_ ∈ [0, 1],

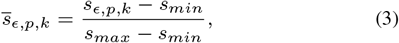

with *s*_*min*_ and *s*_*max*_ being respectively the smallest and the highest sizes of all computed *mvc*. Finally we compute *w*_*ϵ,p,k*_, the *mvc*_*ϵ,p,k*_ score as the geometric mean between (2) and (3),

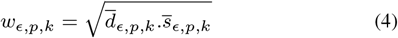

#### 2.1.6 Metavariant species as maximum weighted independent set

The identification of the MVS, can be seen as maximizing the number of non-overlapping *mvc* and their corresponding weight at the same time. This corresponds to a Maximum Weighted Independent Set (MWIS) problem. Several algorithms have been previously proposed by Sakai and colleagues (Sakai *et al.*, 2003). Here, we use the WMIN algorithm to find *MWIS*, the set of all MWISs.

In this context, *MVC* can be represented by a weighted undirected graph *G*(*V, E, W*), where ∀*i, j* ∈ {1*, …,* |*V*|}, *v*_*i*_ ∈ *V* represents *mvc*_*i*_ of weight *w*_*i*_ ∈ *W* and *e*_*ij*_ ∈ *E* ⇔ *mvc*_*i*_ ∩ *mvc*_*j*_ ≠ ∅ and *mvc*_*i*_ ≠ *mvc*_*j*_. We recall here the sketch of the Sakai WMIN algorithm: it takes *G* as input and iterates until *G* = ∅. At each iteration, the following steps are performed (i) detection of the connected components (cc), (ii) in each cc, finding the node that is the maximum weighted independent set, *v*_*i*_ = *mwis* ∈ *MWIS*, if *f* (*v*_*i*_) = *argmax*(*f*), with 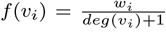 and with *deg*(*v_i_*) the *v*_*i*_ degree. (iii) in each cc, deletion the neighbors of the *mwis* from *G*, and storage of *mwis* in *MWIS* and deletion *mwis*. The fact that this algorithm needs *w*_*i*_ ∈ ℝ* justifies (2) and (3).

**Algorithm 1:**
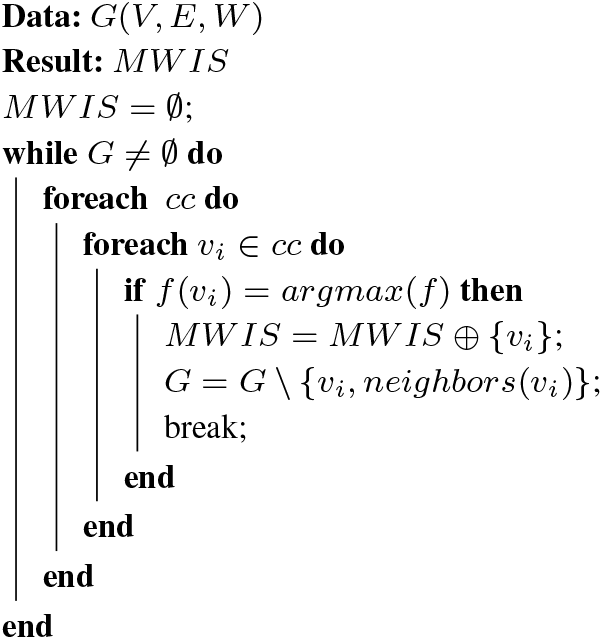
WMIN algorithm from Sakai et *al*.

#### 2.1.7 Selection of metavariants clusters as metavariant species

A metavariant cluster is a potential MVS if it satisfies four criteria applied in the following order: (i) the metavariant cluster is a maximum weighted independent set. (ii) The metavariant cluster occurs in more than *k*_*min*_ populations (set to 4 by default) and corresponding loci have a median depth of coverage higher than *c*_*m*_ (set to 8 by default). (iii) its filtered variable loci have a depth of coverage within *c*_*min*_ = *c*_*m*_ − 2 * *sd*, *c*_*min*_ ≥ 8 by default and *c*_*max*_ ≤ *c*_*m*_ + 2 * *sd* in all samples. (iv) the metavariant cluster contains more than *m_min_*_2_ metavariants (set to 100 by default). More formally,

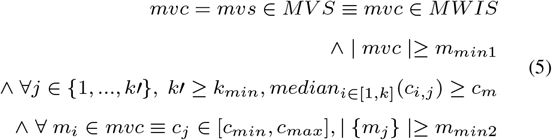

#### 2.1.8 MVS-based population genomic analysis

The allele frequencies of metavariant species are used to compute classical population genomic metrics. This includes the global *F*_*ST*_ according the Wright’s definition (Wright, 1950) such as,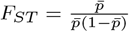, with *p̅* being the mean allele frequency across all samples. We also computed the *LK*, a normalized *F*_*ST*_, such as 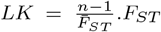, with *F̅*_*ST*_ being the mean *F*_*ST*_ across all loci. *LK* is expected to follow a *χ*^2^ distribution when a large majority of the polymorphic loci are under neutral evolution Lewontin and Krakauer (1973). To estimate the genomic differentiation between MVS populations, we compute the pairwise-*F*_*ST*_ between the different populations (i.e samples).

### 2.2 Implementation of the *mDBSCAN-WMIN* algorithm in *metaVaR*

The metavariants preprocessing step is performed by running *metaVarFilter.pl*, that produces the depth of coverage and frequency matrices from a reference-free vcf file. The *MetaVaR* package was written in R and provides three main functions for MVS construction:
 
1. *tryParam* creates metavariant clusters using several *e, p* values. We used the R package *fitdistrplus* to obtain the log-likelihood of the coverage distribution.
2. *getMWIS* identify the maximum weighted independent sets.
3. *mvc2mvs* applies filters described in (5) to select the MVS and performs population genomic analysis.

metaVaR is available at https://github.com/madoui/metaVaR.

### 2.3 Metavariants set from simulated metagenomic data

We downloaded six bacterial genomes from NCBI (*Escherichia coli* NC_000913.3, *Pseudomonas aeruginosa* NC_002516.2, *Yersinia pestis* NC_003143.1, *Rhizobium tropici* NC_020059.1, *Rhizobium phaseoli* NZ_CP013532.1, *Rhodobacter capsulatus* NC_014034.1). For each genome we created a derived genome that contains 1% of randomly distributed SNPs for *E. coli* and 2% for the others bacteria. We used metaSim (Richter *et al.*, 2008) to simulate Illumina paired-end 100 bp reads from 300bp genomic fragments on seven populations that contained various abundance (Figure 2.A) and various proportion of original and derived genomes (Figure 2.B).

**Fig. 2.**
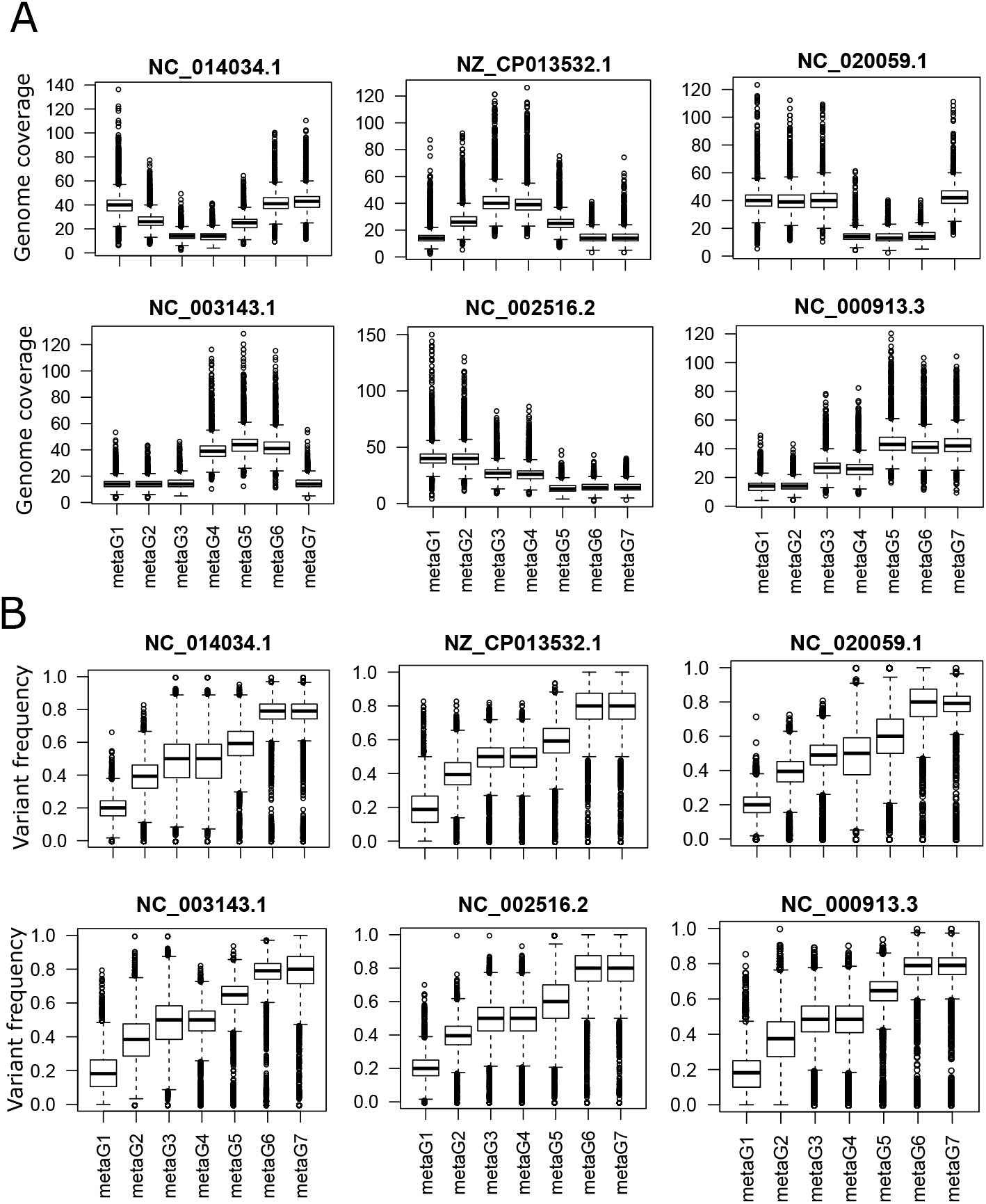
Simulated seven metagenomic dataset on an admixture of six bacterial species containing within-species single nucleotide polymorphism. A. Genome coverage distribution of the species. B. Within-species variants frequencies distribution

To generate the metavariants, *DiscoSNP++* was ran on the total read set with parameter *-b 1*. As a control, the metavariants were relocated on the six original genomes using the -G option. From the VCF file produced by *DiscoSNP++*, the depth of coverage of the biallelic loci and allele frequencies were calculated using *metaVarFilter.pl* with parameters *-a 10 -b 500 -c 7* to keep only loci present in the seven samples.

### 2.4 Metavariants set from real metagenomic data

To test the *mDBSCAN-WMIN* algorithm performances on real metagenomic data, we used metagenomic reads from five marine samples collected in the Mediterranean Sea (Table 1). In a previous study, the reads were processed by *DiscoSNP++*, the metavariants were aligned on the *Oithona nana* genome and the population genomics was performed to estimate the genomic differentiation between samples by pairwise-*F*_*ST*_ and to identify loci under selection (Arif *et al.*, 2018).

**Table 1.** Metavariants clustering performances on simulated metagenomic data. *The computation was performed on Intel(R) Core(TM) i5-7200U CPU

| Algorithm | Software | Recall | Precision | Signal-To-Noise | Purity | Entropy | Time CPU (min)* |
| --- | --- | --- | --- | --- | --- | --- | --- |
| mDBSCAN-WMIN | metaVaR | 0.5523 | **0.9996** | **1691.18** | **0.9999** | **0.008** | 3.96 |
| CLR-kmeans | coseq | **0.5945** | 0.9941 | 99.79 | 0.993 | 0.22 | **1.66** |
| arcsin-GMM | coseq | 0.3007 | 0.998 | 158.24 | 0.9915 | 1.77 | 4.41 |
| logit-GMM | coseq | 0.2685 | 0.9993 | 408.6 | 0.9892 | 1.76 | 5.94 |

In the present study, *DiscoSNP++* was run on the five read sets and the vcf output was filtered using *metaVarFilter.pl* with parameters *-c 20 -b 250 -c 3* that produced two files containing the depth of coverage of metavariant loci and the metavariant frequencies matrices. These two files were used as input for *metaVaR* and other algorithms used for benchmarking (see next section for details).

As a control, the metavariants belonging to *O. nana* were identified by mapping the metavariants back on the *O. nana* genome. The *MWIS* corresponding to *O. nana* was used to compare the genomic differentiation (pairwise-*F*_*ST*_) estimated by *metaVaR* to the reference-based approach previously published by (Arif *et al.*, 2018).

In *O. nana*, loci with higher *LK* values that expected (based on *χ*^2^ distribution) were considered under selection for *p* − *value* ≤ 0.05.

### 2.5 Comparison of *mDBSCAN-WMIN* to other sequence abundance-based clustering algorithms

The depth of coverage matrix was used for metavariants clustering by *mDBSCAN-WMIN* using parameters *e* = (3, 4, 5, 6, 7) and *p* = (5, 8, 10, 12, 15, 20). No clustering algorithm has been explicitly developed for metavariant clustering, however this clustering problem can be solved by other sequence abundance-based clustering algorithms currently used in RNA-seq data analysis to identify co-expressed genes (Rau and Maugis-Rabusseau, 2018). State-of-the-art clustering algorithms has been implemented in *coseq* and tested: (i) centered log-ratio transform and k-means clustering (with k values from 2 to 12), (ii) arcsin transform and Gaussian Mixture Model with same k values as (i), and (iii) logit transform and GMM with same k values as (i).

To evaluate the performances of each clustering algorithm on simulated data, first the clusters were assigned to one single original genome based on the highest proportion of metavariant originated from the same genome. For a cluster, we defined the true positives *TP* as the number of metavariants of the cluster coming from the original genome, the false positives *FP* as the number of metavariants of the cluster not coming from the original genome, the false negatives *FN* the number of metavariants of the original genome not present in the cluster and the true negatives *TN* as the number of metavariants not present in the cluster and the original genome. We computed the recall, precision, signal to noise (STN) of each cluster, as follow 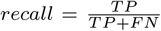, 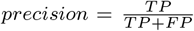, 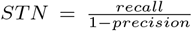, with *TP*, the number true positives, *FN* the number of false negative and *FP* the number of false positive.

The global purity and entropy of the clustering was computed as follow, 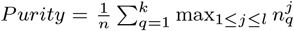 with *n* the number of metavariants, 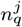 the number of metavariants in cluster *q* that belongs to original species *j* and 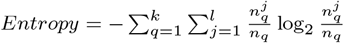 with *n* is the total number of metavariants, *nq* is the total number of samples in cluster *q* and 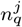 is the number of samples in cluster *q* that belongs to original species *j*.

## 3 Results

### 3.1 Metavariant species as a new modelling of organisms from metagenomic data

In the absence of reference genome to guide metagenomic data analyses for population genomics, we model species only by their variable loci and associated depth of coverage and variant frequencies in different environment samples. We called this model *metavariant species* or MVS and we propose a method to construct them from multisample raw metagenomic data (Figure 1.A). The method is based on the reference-free variant calling from metagenomic reads from different samples performed by *DiscoSNP++* (Figure 1.B).

In the context of metagenomics, the variants are called metavariants and clustered into MVSs. The metavariants are clustered by multiple density-based clustering (*mDBSCAN*) based on the covariation of the depth of coverage of the variable loci across samples (Figure 1.C). Then, the clusters are scored statistically based on the expected depth of coverage of the variable loci in each sample (Figure 1.D). The best clusters are selected by a maximum weighted independent set (WMIN) algorithm (Figure 1.E) and variable loci with a certain minimal coverage are selected to obtain the final MVSs (Figure 1.F). The method was implemented in a *R* package called *metaVaR* that allows users to build and manipulate MVSs for population genomic analyses (Figure 1.G).

### 3.2 Metavariant species on simulated metagenomic data

To test the relevance of MVSs for population genomic analyses, we simulated seven metagenomic data set composed of illumina short reads from an admixture of six bacteria in various abundances (Figure 2.A). Each bacterial species was composed of two strains in various abundance (Figure 2.B). Metavariants were detected by *DiscoSNP++* and filtered resulting in 90,593 metavariants from which the depths of coverage of the variable loci and metavariants frequency matrices were computed.

The metavariants were clustered into MVS candidates using the *mDBSCAN-WMIN* algorithm and the clustering performances were compared to three state-of-the-art algorithms (Rau and Maugis-Rabusseau, 2018): (i) centered log-ratio and kmeans (CLR-kmeans), (ii) arcsin transform + Gaussian mixture model (arcsin-GMM), (iii) logit transform + Gaussian mixture model (logit-GMM).

The clustering performances of the four algorithms are summarised on Table1 and are illustrated on Figure 3. Overall, the *mDBSCAN-WMIN* algorithm obtained the highest precision, signal to noise ratio (STN), purity and entropy. *mDBSCAN-WMIN* is slightly but not significantly less sensitive than *CLR-kmeans* (Figure 3.A), (paired U-test, *P* ≥ 0.05). *mDBSCAN-WMIN* is significantly more precise than the three other algorithms (Figure 3.B) and has a significant higher STN (Figure 3.C) (paired U-test, *P* ≤ 0.05). Moreover, three of the *mDBSCAN-WMIN* clusters over the six contained zero false positives. The arcsin-GMM and logit-GMM methods showed significantly lower recall (paired U-test, *P* ≤ 0.05).

**Fig. 3.**
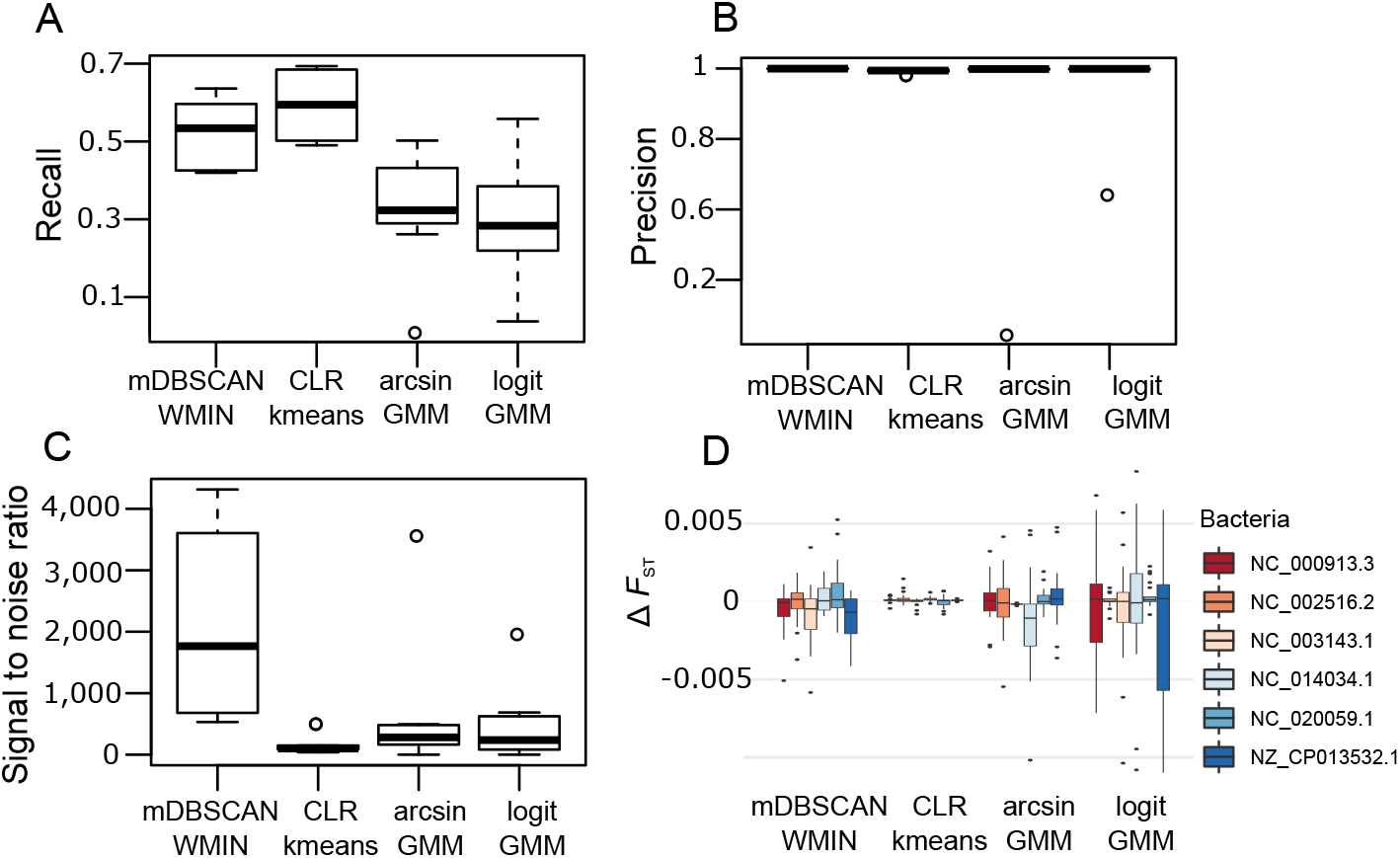
Metavariants clustering performances of four clustering algorithms. A. Recall of metavariants clusters. B. Precision of metavariant clusters. C. Signal to noise of metavariant clusters. D. *F*_*ST*_ difference between MVS and the alignment-based method

The six clusters selected by *mDBSCAN-WMIN* correspond to three different *DBSCAN* parameter settings. Four clusters generated with *e* = 6*, p* = 8 were selected, one cluster for *e* = 6*, p* = 5 and one for *e* = 7*, p* = 12. This illustrates well the fact that according to our scoring method there is not a single (*e, p*) couple to obtain the highest scored metavariants clusters.

The accuracy of the pairwise-*F*_*ST*_ estimation based on MVS was evaluated for the four clustering algorithms and compared to the one obtained by metavariants alignment on the six bacterial genomes (Figure 3.D). Pairwise-*F*_*ST*_ estimated on clusters from the four algorithms showed negligible differences to the reference-based approach with Δ*F*_*ST*_ ≤ 0.01, however the *CLR-kmeans* algorithm showed the lowest Δ*F*_*ST*_ values.

### 3.3 Metavariant species on real metagenomic data

To evaluate the relevance of MVS on real metagenomic data generated from environmental samples containing more complex genomes than the bacterial ones, we selected five metagenomic marine samples known to contain the zooplankter *Oithona nana* in sufficient abundance for population genomic analyses (Arif *et al.*, 2018).

We ran *DiscoSNP++* on the raw data and generated 11.5e6 metavariants filtered into 138,676 metavariants. The metavariants were clustered into MVS using the same four clustering algorithms as previously tested for simulated data. MVSs corresponding to *O. nana* were identified by aligning the metavariant on its genome. *mDBSCAN-WMIN* and *CLR-kmeans* detected the *O. nana* MVS, but the two other methods generated the maximal number of MVS allowed by the parameters (i.e 12 clusters) with no clusters assigned to *O. nana*, thus these latter is not considered further (Table 2). The *O. nana* MVS built by *mDBSCAN-WMIN* is less complete but more accurate than the one built by *CLR-kmeans* (Table 2). The metavariants that have not been relocated on the *O. nana* may include missing part of the genome assembly.

**Table 2.** Metavariants clustering performances on real metagenomic data. *Performances for the O. nana cluster. Time CPU is in minutes.

| Algorithm | Software | Number of cluster | Recall* | Precision* | Signal-To-Noise* | Time CPU |
| --- | --- | --- | --- | --- | --- | --- |
| mDBSCAN-WMIN | metaVaR | 3 | 0.1592 | **0.8144** | **1691.18** | **1.48** |
| CLR-kmeans | coseq | 4 | **0.1662** | 0.7182 | 99.79 | 4.5 |
| arcsin-GMM | coseq | 12 | - | - | - | 5.53 |
| logit-GMM | coseq | 12 | - | - | - | 4.9 |

The pairwise-*F*_*ST*_ matrices of the *O. nana* MVSs (Figure 4.A) showed small differences compared to alignment-based *F*_*ST*_ values (Δ*F*_*ST*_ ≤ 0.03) (Figure 4.B). To illustrate potential MVS applications, several down-stream analyses were performed including isolation-by-distance (IBD) (Figure 4.C), species co-differentiation (Figure 4.D) and natural selection tests (Figure 4.E-F). In the Mediterranean Sea, the Lagrangian distances between the Western and Eastern Basins sampling sites (S10, 11, 12 and S24, 26 respectively) explain significantly the *O. nana* genomic differentiation (Mantel *r* = 0.73, *p* − *value* ≤ 0.05) (Figure 1.C) Mantel (1967). The comparison of the genomic differentiation trends between three MVSs (detected by the *mDBSCAN-WMIN*) showed a negative correlation between MVS2 and MVS3 but no co-differentiation patterns between other MVS pairs.

**Fig. 4.**
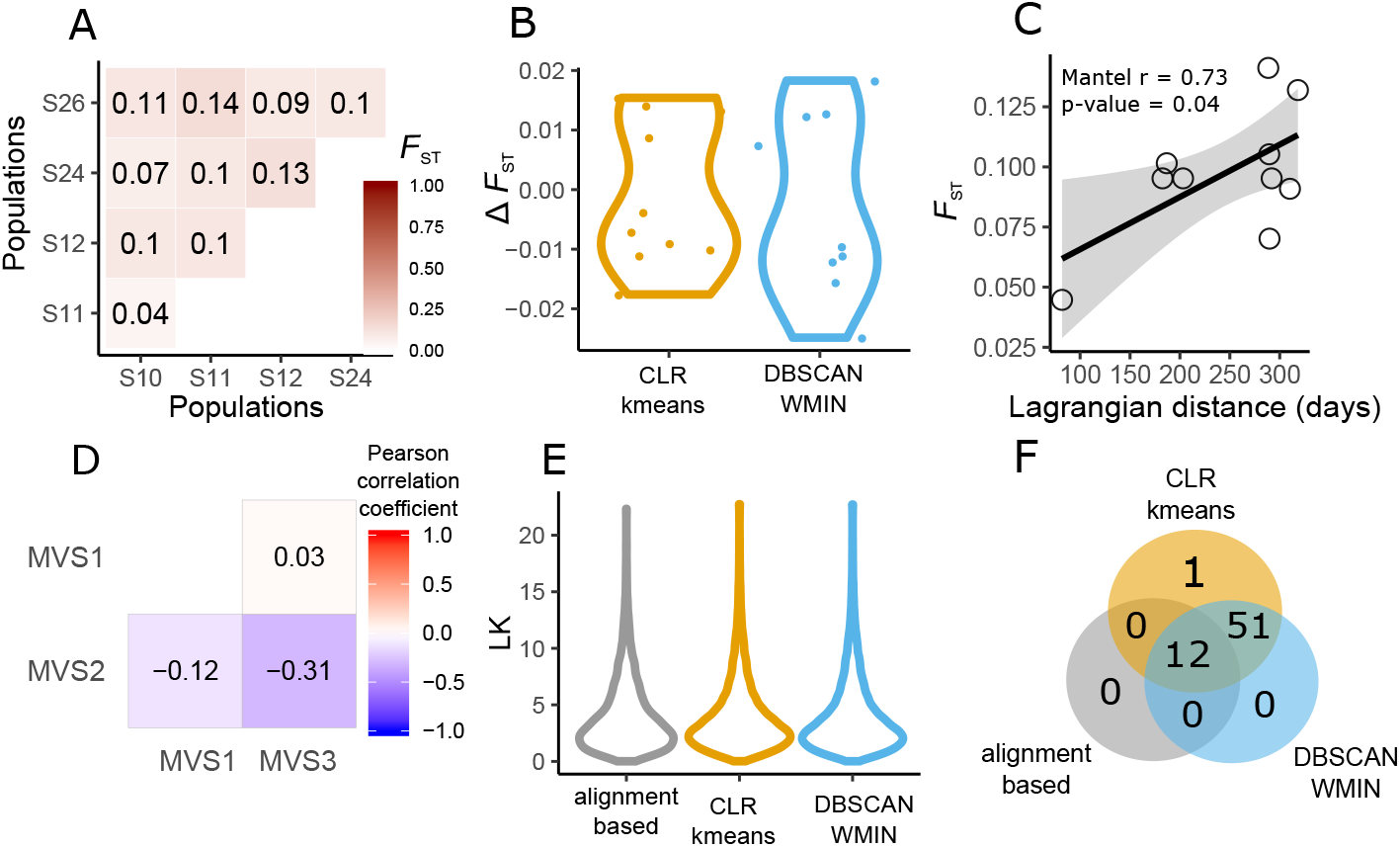
Examples of applications and accuracy of metavariants species. A. Genomic differentiation based on pairwise *F*_*ST*_ of O. nana populations from the MVS built with DBSCAN-WMIN. . B. Difference of *F*_*ST*_ values between alignment-based method and MVSs. C. Mantel test with Lagrangian distances. D. Co-differentiation between species. E. LK distribution. F. Venn diagram of loci predicted under selection (LK p-value ≤ 0.001) by the different approaches.

Loci under selection in Mediterranean populations of *O. nana* were identified based on LK outlier values produced by the MVS-based and the alignment-based methods. For the three approaches the LK distribution supports neutral evolution of most of the *O. nana* polymorphic sites (Figure 4.E). The prediction of loci under selection based on the extreme LK values by the three approaches showed that all loci predicted by the alignment-based approach are found in the *O. nana* MVSs and more loci are found under selection with MVSs where both clustering methods differs only form one loci (Figure 4.F).

## 4 Discussion

### 4.1 Modeling species nucleotide polymorphism by metavariant species

Large-scale nucleotide polymorphism detection requires traditionally reference sequences like genome or transcriptome assemblies. Such resources are lacking for most of species creating a bottleneck in population genomic investigations by drastically reducing the choice of studied species among the sequenced ones. Unfortunately, the set of species with a transcriptome or genome reference does not well represent the whole genomic landscape complexity of small eukaryote-rich biomes present notably in the oceans (Carradec *et al.*, 2018). The MVS representation allows to bypass this barrier and to investigate a much larger number of species including unknown species for which poor or no genomic resources are available.

However, the MVS modelling requires a minimal amount of genomic information that includes the variable loci of single copy region and their variant frequencies in the different samples. The information is sufficient to perform basic population genomic analysis like genomic differentiation and detection of loci under natural selection. The extraction of this information from raw metagenomic data through the use of *DiscoSNP++* does not need reference or assembly and generates accurate variant frequencies in a reasonable time and computational resources even for very large dataset (Arif *et al.*, 2018).

To provide an ecological insight out of the MVSs, we recommend to perform a taxonomical assignation of the MVSs. This is feasible using the variable loci sequences generated by *DiscoSNP++* and to align them to public sequence databases. Due to the short length of the loci, this task can be performed using classical short reads aligners.

### 4.2 The challenge of metavariants classification by species

The accuracy of the genomic differentiation estimation depends on the sample size and the number of markers. Having an exhaustive SNPs set is not mandatory but a large set (>1000 SNPs) is preferable (Willing *et al.*, 2012). Thus, the number of metavariants in a MVS to consider it as valid is critical. Moreover, loci under selection represents very often a small fraction of the genome, the increase of the metavariant number in a MVS helps to detect such loci but it remains crucial to avoid false positives that will generate biased *F*_*ST*_ values. In the metagenomic context, the false positives are metavariants assigned to a species they do not belong to. *F*_*ST*_ values derived from mis-assigned metavariants may generate outliers and support false natural selection. In this context, one can consider the precision as the first criteria to prioritize. Then, the clustering of the metavariants by *mDBSCAN-WMIN* and *CLR-kmeans* gave the best clustering results on simulated and real data. While the first method is more specific but less sensitive, these two methods have also their own limits. *mDBSCAN-WMIN* needs the try of multiple clustering parameters (*e, p*), several tries might be necessary to obtain all possible MVSs and *CLR-kmeans* needs the try of multiple *k* with no prior knowledge, thus the optimal *k* can be missed. However, *mDBSCAN-WMIN* will not cluster all metavariants

### 4.3 Toward a holistic view of microorganism genomic differentiation and natural selection

Current population genomic analyses focus on one single species at a time. Thanks to MVSs, the genomic differentiation of several species can be modelled simultaneously and hypotheses like isolation-by-distance can be tested on each species. The genomic differentiation of MVSs can be compared and species sharing common differentiation profiles can be identified to illustrate possible co-differentiation or similar gene flow.

Another relevant MVSs application concerns natural selection. The ratio of loci under selection over the total number of variable loci is an interesting metric to estimate the force of the natural selection on the species molecular evolution. This ratio can be computed on each MVS and compared to assess the relative effect of natural selection.

### 4.4 Current limits and future developments for metavariant species-based population genomics

Pairwise-*F*_*ST*_ remains currently a robust metric to draw the silhouette of the genomic differentiation from metagenomic data. The absence of genotypes and haplotypes and their relative frequencies disables intra-population analysis and does not allow to compute *F*_*ST*_ p-value. Moreover, the use of population genomic tools enabling the estimation of nucleotide diversity, the identification of genomic structure and the test of evolutionary trajectories and past demographic events is not yet possible. For these reasons, future developments focusing on variant phasing and haplotyping from metagenomic data will greatly help to improve MVSs application. In this context, the use of long read sequencing technologies will be a strong advantage by supporting long-range haplotypes spanning several kilobases.

## Conclusion

Thanks to MVS, the population genomic analyses of unknown organisms is now feasible without reference genome or genome assembly. MVSs are suitable for genomic differentiation and natural selection analysis. The simultaneous access to several species nucleotide polymorphisms present in the same ecosystem allows for a holistic view of microorganism genomic differentiation and adaptation. Future development will try to reconstruct the species haplotypes based on metavariant species and will allow a more accurate view of species evolution.

## Acknowledgements

We acknowledge David Vallenet, Guillaume Gautreau and Adelme Bazin for helpful discussions.

## Funding

This work was supported by the Commissariat à l’Energie Atomique et aux Energies Alternatives (CEA) and by IRISA-Inria.

